# Topological Closure Drives Structural Stabilization and Fast Cooperative Dynamics in Crowded Circular Polysomes

**DOI:** 10.64898/2026.08.31.748270

**Authors:** Hideki Kobayashi, Horacio V. Guzman

## Abstract

In linear polysomes, excluded-volume interactions among ribosomes can induce dimensional reduction of mRNA. Yet linear architectures allow steric stress to relax at open ends– limiting how strongly crowding can remodel the mRNA’s structure and dynamics. Using coarse-grained molecular-dynamics simulations, we compare circular and linear polysomes over a range of ribosome densities. Circular closure selects a predominantly quasi-planar global conformational ensemble, as indicated by a shape dimensionality *d*_shape_ ≈ 2 over a range of ribosome densities. Crucially, circular topology and ribosome crowding act cooperatively to suppress structural fluctuations. While closure alone or linear crowding reduces relative global size fluctuations (Δ*R*_g_/*R*_g_) only to ≈ 0.16, their combined effect drives this fluctuation down to ≈ 0.07. Within this stabilized architecture, increasing ribosome density drives a distinct in-plane reorganization: the ring becomes more isotropic, global size fluctuations are strongly suppressed, and the scaling exponent increases toward *ν*≃ 0.74–0.77, consistent with two-dimensional self-avoiding walk-like value over the accessible finite-size window, 1000 ≤*N*≤ 4969. Closure shortens the radius-of-gyration decorrelation time of circular polysomes by 40-fold relative to matched linear systems, reflecting the topological elimination of free ends. Within this closure-selected ensemble, ribosome crowding further reduces the decorrelation time by up to 20% at the highest density. A fluctuation-informed crossover model links the density dependence of the global scaling exponent to inter-ribosomal subchain statistics. These results distinguish the geometric role of circular closure from the density-dependent steric response that it enables, revealing a confined yet dynamically responsive conformational regime for circular polysomes.

## I. INTRODUCTION

Polysomes, as multiple ribosomes translating a single mRNA, are known to adopt diverse spatial organizations^1–9^ in cells, including circular topologies. In eukaryotic translation, such circularization has long been discussed in connection with enhanced translational efficiency, because the proximity of the initiation and termination regions is thought to facilitate ribosome recycling and reinitiation^10–12^. At the same time, however, a higher local density of ribosomes along a circular polysome may impose strong steric constraints on the mRNA backbone, potentially leading to enhanced mechanical strain, translational traffic, or ribosome collisions^13–16^. Thus, whether circular topology merely promotes recycling or also generates a mechanically and conformationally favorable polysome state remains an open biophysical question.

In polymer physics, the conformation of ring polymers is strongly influenced by stiffness and self-avoidance. For semiflexible rings, increasing the persistence length drives a crossover from flexible three-dimensional conformations toward more planar and loop-like shapes^17,18^. Polysomes differ from such passive rings in an essential way: the polymer backbone is decorated by ribosomes, whose finite size and steric repulsion impose additional geometric constraints on the accessible configuration space. Consequently, conformational restriction in polysomes need not arise from chain stiffening alone, but may result from a self-induced reduction of the effective dimensionality driven by ribosome crowding.

In our previous multiscale simulations of linear polysomes, we demonstrated that bulky ribosomes sandwiching the mRNA backbone create an anisotropic (*i*.*e*. arising from the size difference of the small and large ribosomal units), sheetlike “steric corridor”^19^. Along the longitudinal axis, large ribosome diameters impose severe steric barriers that suppress axial fluctuations. Conversely, lateral fluctuations remain relatively unconstrained because the mRNA exits near the peripheral edge where the steric obstacle is thinner. While linear topologies allow these anisotropic constraints to relax at terminal free ends, circular polysomes eliminate boundaries, enforcing synergistic topological and crowding constraints. The central question is whether topological closure merely removes end effects or generates an effectively distinct, sterically constrained regime with altered scaling and dynamics at biologically relevant mRNA length scales^20^.

Here, we address this question by performing large-scale coarse-grained simulations of circular and linear polysomes decorated with ribosomes over a range of loading densities. We analyze the conformational ensemble using the radius of gyration, scaling exponents, shape dimensionality, prolateness, and relaxation dynamics. We find that circular topology redistributes steric constraints along the backbone and yields a more restricted conformational ensemble than the linear case. In particular, circular polysomes exhibit reduced fluctuations, modified scaling behavior, and distinct relaxation characteristics, consistent with strongly constrained but still dynamically accessible conformational-states. These results demonstrate that topological closure and ribosome crowding together define a distinct conformational regime in polysomes within biologically relevant length scales.

## II. MODEL AND NUMERICAL METHOD

To simulate the structural and spatial dynamics of translationally active polysomes, we construct a coarse-grained molecular dynamics model representing an mRNA transcript bound to multiple ribosomes.

### Coarse-Grained Polymer Model for mRNA

The mRNA backbone is modeled as a coarse-grained flexible polymer using a modified Kremer–Grest model^21^. Each nucleotide unit is represented by a single monomer with an effective diameter *σ*_m_ = *u* = 0.6 nm, which serves as the fundamental spatial unit, and all physical quantities are expressed in reduced units based on this length. The energy unit is *k*_*B*_*T*, the mass unit *m* is the mass of a monomer and the unit time *τ* correspond to 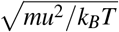. Consecutive monomers along the chain are linked via a Finitely Extensible Nonlinear Elastic (FENE) potential:

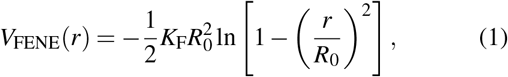

where *K*_F_ = 30 *k*_B_*T* /*u*^2^ denotes the spring constant and *R*_0_ = 1.5*u* defines the maximum allowable bond length.

Non-bonded monomer-monomer excluded volume interactions are captured by a purely repulsive Weeks-Chandler-Andersen (WCA) potential:

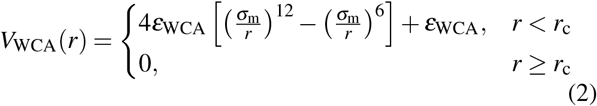

with interaction strength *ε*_WCA_ = 1.0 *k*_B_*T* and a potential cutoff at *r*_c_ = 2^1/6^*σ*_m_. We evaluate chain length dependencies using total monomer counts of *N* = 1000, 2000, 3000, 4060, and 4969.

In addition to intrachain elasticity, local chain stiffness is controlled by a harmonic angle potential applied across three consecutive monomers:

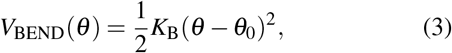

where *K*_B_ = 50.0 *k*_B_*T* /rad^2^. We set the equilibrium bending angle to either *θ*_0_ = *π*/2 or 2*π*/3, consistent with parameterizations calibrated to reproduce experimental end-to-end distances of cellular mRNAs^22^.

To model circular polysomes, an end-to-end loop closure constraint is introduced by connecting the initial monomer (*i* = 1) and final monomer (*i* = *N*) of the mRNA chain via a harmonic tethering potential:

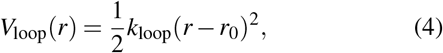

where the spring constant is set to *k*_loop_ = 50 *k*_B_*T* /*u*^2^ and the equilibrium terminal distance is fixed at *r*_0_ = 1.2*u*. This harmonic constraint mimics the non-covalent, protein-mediated bridging (e.g., via initiation factor complexes) between the 5’ and 3’ ends rather than a continuous phosphodiester backbone. Accordingly, the bending potential (Eq. 3) is applied exclusively to internal mRNA triplets, allowing localized conformational flexibility at the junction while strictly enforcing topological closure.

### Ribosome Architecture and mRNA Coupling

Ribosomes are explicitly incorporated as two contacting spheres representing the small (SSU) and large (LSU) subunits, with diameters *σ*_S_ = 20*u* (12 nm) and *σ*_L_ = 30*u* (18 nm), respectively. The geometric attachment of ribosomes to mRNA is maintained through specific footprint tethers: each ribosome spans a dedicated 30-monomer mRNA segment. These 30 footprint beads are harmonically anchored to the SSU center using:

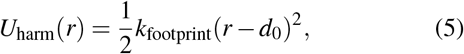

with *k*_footprint_ = 10 *k*_B_*T* /*u*^2^ and equilibrium separation *d*_0_ = 10*u*. Furthermore, the midpoint monomer of this 30-bead footprint is simultaneously tethered to the LSU center via the identical potential (*k*_footprint_ = 10 *k*_B_*T* /*u*^2^, *d*_0_ = 10*u*). This dual attachment sandwiches the mRNA backbone between the subunits, establishing an effective ribosome long-axis span of 45*u* (27 nm) while preserving a realistic 18 nm lateral thickness. Ribosomes are periodically loaded onto the transcript with defined inter-ribosomal gaps of 30 × *N*_space_ monomers.

### Steric Interactions and Numerical Implementation

Inter-ribosomal steric hindrance and ribosome-monomer interactions are modeled via a smoothed Gaussian Core Potential (GCP):

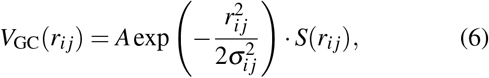

where *A* = 5 *k*_B_*T* for ribosome-ribosome pairs and *A* = 10 *k*_B_*T* for ribosome–monomer pairs. The effective interaction radius is given by *σ*_*i j*_ = (*r*_*i*_ + *r* _*j*_)/2. To ensure continuous derivatives and numerical stability, the potential is modulated by a smooth switching function *S*(*r*_*i j*_):

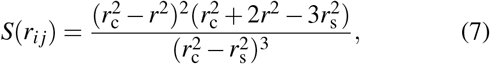

active between *r*_s_ = 1.1*σ*_*i j*_ and *r*_c_ = 1.2*σ*_*i j*_.

Simulations are executed in the NVT ensemble using HOOMD-blue^23^ with a Langevin thermostat (*k*_B_*T* = 1.0, friction coefficient Γ = 0.1, integration timestep *dt* = 0.005*τ*).

Neighbor searching is accelerated using a hierarchical tree-based neighbor list algorithm^24,25^, mitigating memory constraints associated with cell lists in large systems. To prevent inter-molecular artifacts, 32 non-interacting polysomes are simulated simultaneously, each placed within an independent cubic box of volume (2000*u*)^3^, ensuring dilute regime conditions.

## III. RESULTS

### A. The estimation of equilibration time

To establish the appropriate equilibration time for our simulations, we evaluate the characteristic relaxation time of the circular polysomes. This is achieved by calculating the time-displaced auto-correlation function of the fluctuation of the radius of gyration, ⟨*δ R*_g_(*t*)*δ R*_g_(0)⟩, where *δ R*_g_(*t*) = *R*_g_(*t*) − ⟨*R*_g_⟩, for *N* = 1000 across various *N*_space_ values. The squared radius of gyration is defined as

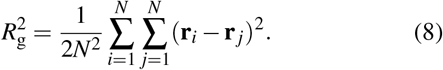

We approximate this auto-correlation function using a stretched exponential decay,

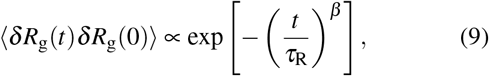

in order to capture the coexistence of multiple dynamical modes within the circular polysome. A stretching exponent *β* < 1 indicates a broad distribution of relaxation times resulting from complex, multi-modal dynamics. The effective decorrelation time *τ*_dec_ is then estimated by

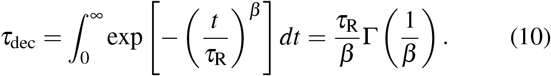

As shown in Fig. 2(left), the stretched exponential fitting accurately describes the relaxation of circular polysomes. To investigate the density dependence of ⟨ *δ R*_g_(*t*)*δ R*_g_(0) ⟩, it is instructive to compare the real-ribosome system with the ghost-ribosome condition. As shown in Table I, in the ghost circular system, although *τ*_dec_ appears to increase slightly with decreasing *N*_space_, this variation remains within a narrow baseline of ∼ 8% across all *N*_space_, while *β* shows no systematic trend and exhibits only minor fluctuations. By contrast, the real-ribosome circular system exhibits a distinct reduction of approximately 20% in *τ*_dec_ at the highest ribosome density (*N*_space_ = 5) relative to the lowest ribosome density (*N*_space_ = 35). Because this reduction clearly exceeds the baseline from the ghost-ribosome cases, it represents a genuine systematic trend: dense steric corridors in a closed topology suppress complex local fluctuations—as also reflected by the exponent *β* increasing toward unity—and force the system to relax more cooperatively and rapidly. Crucially, for the real-ribosome system at *N*_space_ = 5, the decorrelation time *τ*_dec_ ≈ 9.2 × 10^2^*τ* ≈ 0.01Γ*N*^2^*τ* is close to the theoretical Rouse time of an unconstrained passive ring (*τ*_Rouse,ring_ ≈ 0.0085Γ*N*^2^*τ*).

**FIG. 1.**
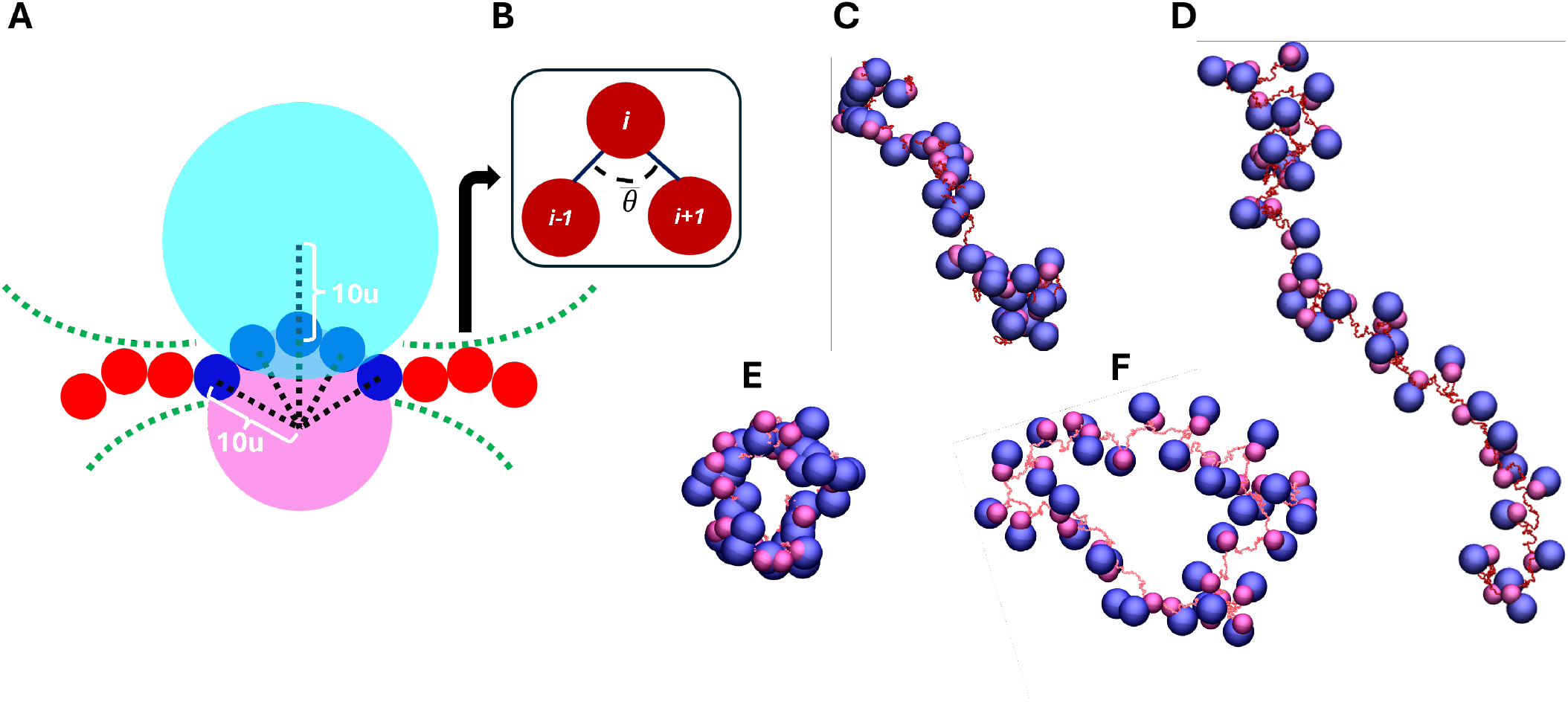
(A) Schematic illustration of the coarse-grained ribosome-mRNA model. The ribosome consists of a large subunit (cyan) and a small subunit (pink). The mRNA backbone is represented by a chain of monomers (red). Within the 30-monomer footprint (blue), monomers are harmonically constrained to the center of the small subunit (black dashed lines) to simulate surface attachment. For clarity, only 5 monomers are illustrated to represent the 30-monomer segment. Additionally, the central monomer of this segment is anchored to the center of the large subunit via a harmonic potential (black dotted line). This dual-anchoring construction ensures that the mRNA is sandwiched between the subunits, effectively forming a single functional unit with an elongated geometry. (B) Schematic illustration of the bending angle definition. Three consecutive monomers, indexed as *i* − 1, *i*, and *i* + 1, define the local configuration of the mRNA backbone. The bending angle *θ* is the angle formed by the two bond vectors, 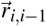 and 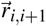, originating from monomer *i*. (C) and (D), show equilibrated mRNA-ribosome conformations for the linear architectures. Both snapshots show a system with *N* = 4, 969, *N*_space_ = 5, and *θ* = *π*/2. (C) In the ghost-ribosome model, the lack of inter-ribosomal excluded volume leads to a compact mRNA configuration. (D) In the real-ribosome case, repulsive interactions induce a structural expansion, resulting in the observed non-SAW scaling behavior. (E) and (F), show equilibrated mRNA-ribosome conformations for the circular architectures. Both snapshots show a system with *N* = 4, 969, *N*_space_ = 5, and *θ* = *π*/2. This makes easier the visual comparison with the linear polysomes from panels (C) and (D). Similarly, the ghost-ribosome model, the lack of inter-ribosomal excluded volume and this leads to a compact mRNA ring conformations (E). While, in the real-ribosome case (F) repulsive interactions induce a structural expansion. Note that in our model, both subunits are treated as repulsive Gaussian units, meaning they exert a purely repulsive force on one another. For illustration purposes we chose the circular polysome snapshots from the cluster with larger *R*_*g*_.

**FIG. 2.**
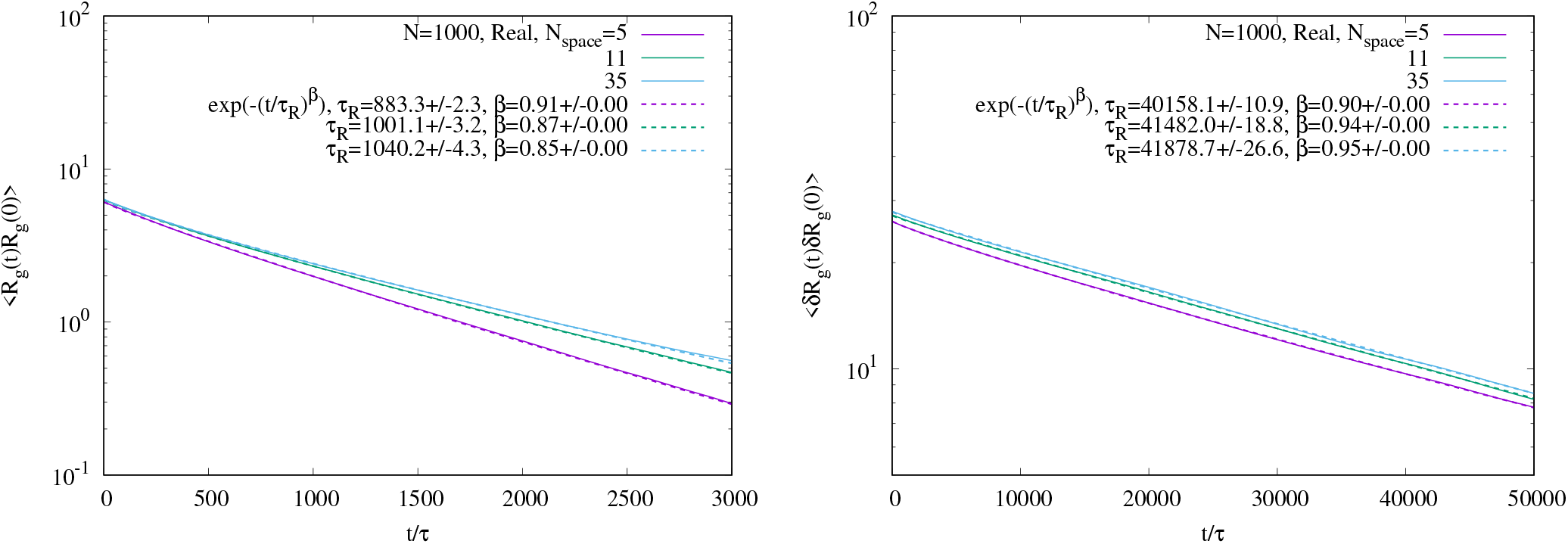
Autocorrelation function of the radius of gyration ⟨*δ R*_g_(*t*)*δ R*_g_(0)⟩ as a function of time *t* for *N* = 1000 monomers. The curves represent ribosome spacings *N*_space_ = 5, 11, and 35, from bottom to top. The dashed line represents least-squares fits to exp (*t*/*τ*_R_) ^β^ . Left: Circular polysome. Right: Linear polysome.

**TABLE I.**
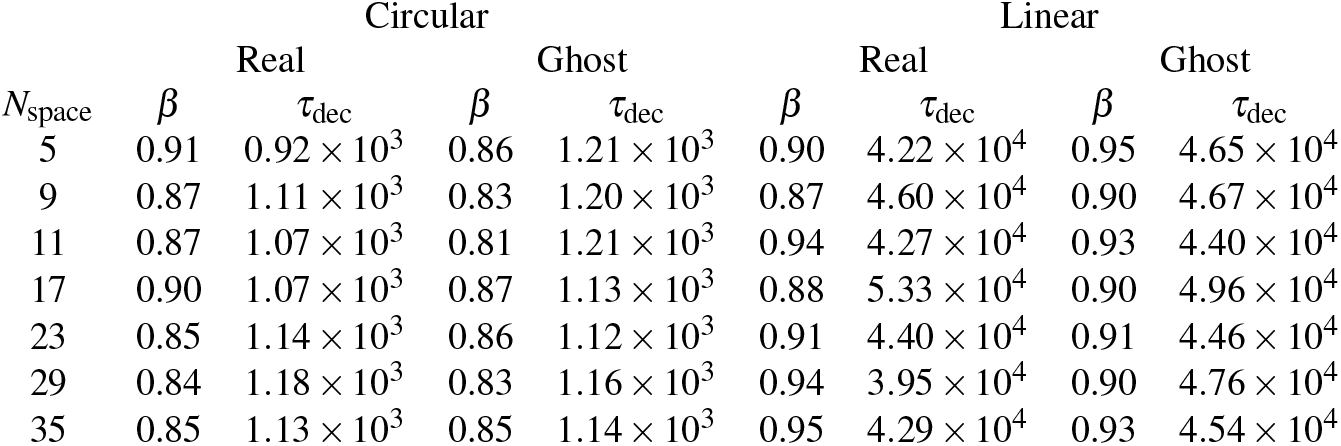
Fitting parameters *β* and effective decorrelation time *τ*_dec_ of the autocorrelation function of *R*_g_ for various ribosome spacing, *N*_space_ with monomer number *N* = 1000. Note that data for the linear polysomes is partly (namely *N*_*space*_ = 5, 9, 11) from previous work^19^

To clarify this mechanism, we also plot ⟨*δ R*_g_(*t*)*δ R*_g_(0) ⟩ for linear polysomes in Fig. 2(right). Applying the same comparative perspective, the ghost linear system exhibits baseline intrinsic fluctuations of ∼ 10% in *τ*_dec_. When steric interactions are introduced in the real linear system, the amplitude of these fluctuations increases (varying broadly between 3.9 × 10^4^ and 5.4 × 10^4^*τ*). However, unlike the circular topology, these amplified fluctuations in the real linear system show no systematic or directional trend with increasing ribosome density (decreasing *N*_space_). Meanwhile, the exponent *β* for real linear polysomes tends to decrease away from unity at higher ribosome densities, indicating that dense loading enhances local complex fluctuations and broadens the relaxation spectrum rather than accelerating global relaxation. Strikingly, the absolute value of *τ*_dec_ for linear polysomes is roughly 48 times slower than that of circular polysomes, regardless of the density. These contrasts demonstrate that the dense steric corridor alone cannot strongly coordinate the global conformational fluctuations. Rather, the cooperative-suppression of dense steric corridors (≈20%) and the circular topology (≈40-fold) suppresses multimodal noise and drives fast, cooperative relaxation dynamics.

The effective decorrelation times, *τ*_dec_, for both linear and circular polysomes with *N* = 1000 across various *N*_space_ are summarized in Table I. We estimate *τ*_dec_(*N*) for arbitrary chain lengths as *α*Γ*N*^2^*τ*, where the prefactor *α* is determined from these characteristic times. Prior to data collection, all systems were equilibrated for a duration of *τ*_eq_ = 3*τ*_dec_ (ensuring that residual correlations decay to exp(−3) ≈ 0.05).

Subsequent production runs were conducted over this equilibrated window using 32 independent replicas to guarantee robust statistical sampling.

### B. Scaling Analysis

The scaling exponent *ν* describes how the spatial size of a polymer chain scales with the number of monomers *N* according to 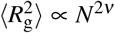. To determine the scaling exponent *ν* for circular polysomes, we calculated the mean-square gyration radius *R*^2^ as a function of chain length *N*, ranging from 1000 to 4969, for various inter-ribosome spacings *N*_space_. In contrast to linear polysomes, circular polysomes lack free ends. Consequently, the steric corridor effect cannot escape through terminal boundaries and is uniformly distributed along the mRNA backbone. As a result, higher values of *ν* are expected at small *N*_space_, whereas systems with large *N*_space_ allow us to evaluate the baseline scaling behaviour in the unconstrained limit.

For *θ* = *π*/2, the scaling exponent *ν* reaches 0.76 ± 0.01 at *N*_space_ = 5 and 0.73 ± 0.01 at *N*_space_ = 11, as shown in Fig. 3(left). Unless otherwise noted, all fitting values are reported as the least-squares fitted value ± fitting uncertainty. Both values for the circular polysome are larger than those observed for linear polysomes, with *ν* ≈ 0.73 and 0.71 for *N*_space_ = 5 and 11, respectively. As *N*_space_ increases to 23 and 35, *ν* decreases to 0.59 ±0.03 and 0.58 ±0.02, respectively, both of which are identical within fitting uncertainty to the standard three-dimensional self-avoiding walk (3D-SAW) value of 0.588. This indicates that the steric corridor effect no longer significantly affects the backbone conformation for *N*_space_ ≥ 23.

**FIG. 3.**
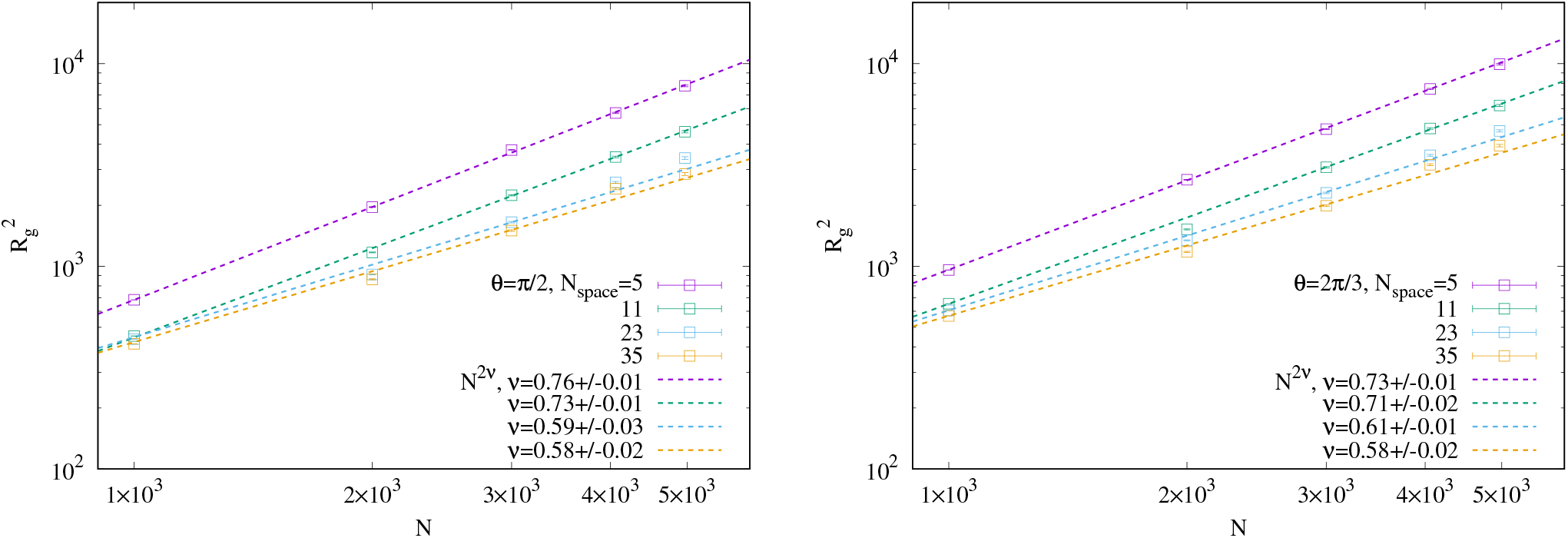
Mean-square radius of gyration, 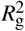, as a function of the number of monomers *N*. The open squares represent ribosome spacings *N*_space_ = 5, 11, and 35, from top to bottom. The dashed lines indicate least-squares fits to the scaling relation 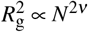. The error bars in the graphs indicate the standard error across 32 independent replicas. Left: The bending angle *θ* = *π*/2. Right: *θ* = 2*π*/3.

As *N*_space_ decreases, the global size scaling of circular polysomes crosses over from a 3D-SAW-like regime toward an exponent close to the 2D-SAW value. This crossover should be distinguished from the global shape dimensionality discussed below: circular closure yields *d*_shape_ ≈ 2 even at low ribosome density, whereas the density dependence of *ν* reports how steric crowding reorganizes conformations within this closure-selected quasi-planar ensemble. Within the Flory-type operational interpretation, the dense-limit exponent corresponds to an effective dimension close to two. We use this correspondence as a convenient description of the observed scaling crossover rather than as a direct measurement of geometric dimensionality. Furthermore, assuming the existence of a characteristic threshold *N*_c_ for the enhancement of *ν*, these data indicate that it falls within the range 11 < *N*_c_ < 23, as *ν* converges to the standard 3D-SAW value for *N*_space_ ≥ 23.

Qualitatively similar behavior is observed for *θ* = 2*π*/3, as shown in Fig. 3(right), yielding *ν* = 0.73 ± 0.01 and 0.71 ± 0.02 for *N*_space_ = 5 and 11, respectively. These slightly reduced values of *ν* compared to the *θ* = *π*/2 case indicate that the increased physical distance between neighboring ribosomes, driven by the longer persistence length, weakens the steric corridor effect. This demonstrates that the self-induced dimensional reduction in polysomes is primarily governed by local ribosomal crowding rather than the intrinsic bending rigidity of the bare mRNA backbone. As *N*_space_ increases to 23, *ν* decreases to 0.61 ± 0.01, which is consistent with the standard 3D-SAW value (0.588) within two standard errors (2*σ*), though the slightly higher mean value may reflect a subtle residual steric effect near the crossover threshold. With further increasing *N*_space_ to 35, *ν* reaches 0.58 ± 0.02, fully converging to the 3D-SAW limit.

Having established that increasing ribosome density enhances the scaling exponent to *ν* ≈ 0.76 (in the quasi-two-dimensional regime), it is crucial to verify whether this structural expansion stems from direct ribosomal collisions or from the steric corridor effect operating consistently across all *N*_space_. To rule out direct collisions as the primary driver of the scaling behavior, we evaluated the probability density function (PDF), *P*(*r*), of the center-to-center distance *r* between neighboring large ribosomal subunits.

As shown in Fig. 4(left), the peak position of *P*(*r*_rib_) shifts toward larger distances *r*_rib_ with increasing *N*_space_, while *P*(*r*_rib_) remains extremely small within the effective interaction range of the large subunit (*r*_rib_ < 36*u*, corresponding to the potential cutoff distance *r*_c_ = 1.2*σ*_L_). Quantitatively, the collision probability between large subunits—calculated by integrating *P*(*r*_rib_) over 0 < *r*_rib_ < 36*u*—is approximately 2% even for the highest density case (*N*_space_ = 5). This demonstrates that direct physical collisions between large ribosomal subunits are rare and play a negligible role in the enhancement of *ν*.

**FIG. 4.**
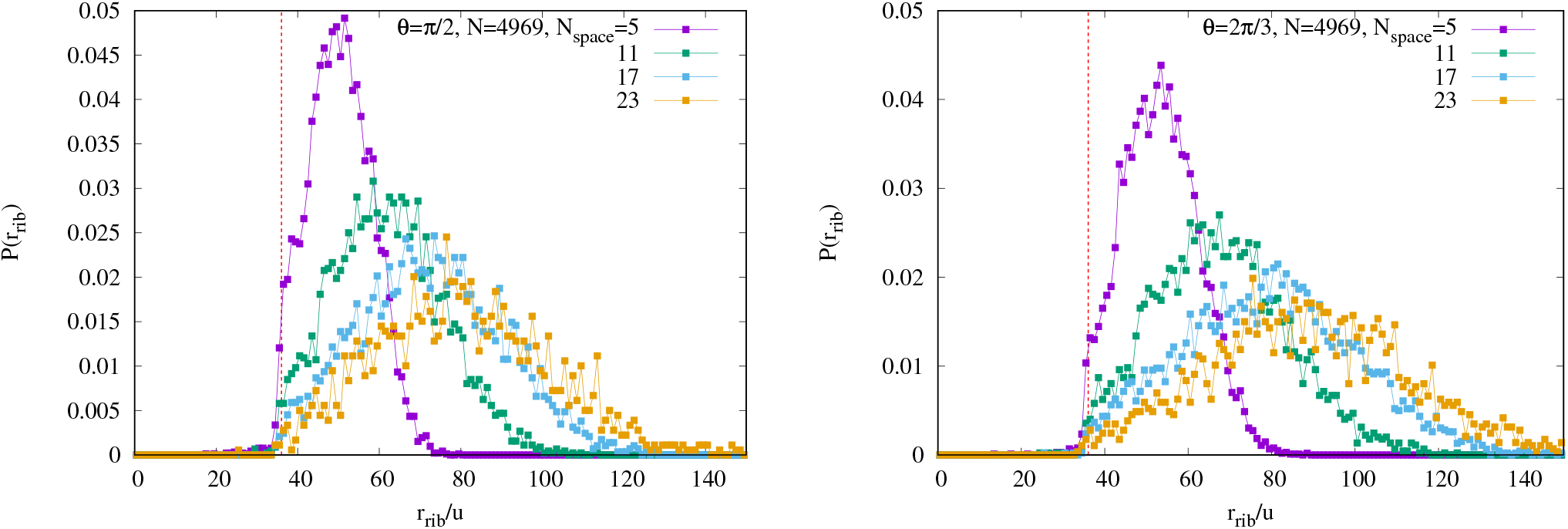
Probability density function of the distance between the large subunits of adjacent ribosomes for monomer number *N* = 4969 at ribosome spacings *N*_space_ = 5, 11, 17, and 23. The peaks shift from left to right with increasing *N*_space_. The vertical dashed line indicate *r*_rib_ = 36*u*. Left: The bending angle *θ* = *π*/2. Right: *θ* = 2*π*/3.

The average inter-ribosomal distances *r*_rib_/*u* are 50.3 ±7.9, 62.1 ±13.4, 72.6 ±17.5, and 75.1 ±20.9 for *N*_space_ = 5, 11, 17, and 23, respectively, where the uncertainties represent the standard deviation (STD). Naturally, all mean values exceed the effective interaction range of the large subunit. The increase in the STD of *r*_rib_ with increasing *N*_space_ implies that the structural fluctuations of the mRNA backbone increase.

The system with *θ* = 2*π*/3 shows qualitatively identical behavior, as shown in Fig. 4(right), although *P*(*r*_rib_) displays a broader distribution with a lower peak height. This broadening reflects the increased structural flexibility of the polysome resulting from the larger spatial separation between adjacent ribosomes. For quantitative comparison, the average inter-ribosomal distances *r*_rib_/*u* are 53.3 ±9.2, 67.6 ±15.7, 78.9 ±20.1, and 81.5± 23.9 for *N*_space_ = 5, 11, 17, and 23, respectively, where the uncertainties also represent STD. These values are systematically larger than those observed for *θ* = *π*/2, confirming that the polysome architecture for *θ* = *π*/2 is distinctly more rigid than that for *θ* = 2*π*/3.

### C. Shape anisotropy

To directly represent the geometric anisotropy of polysomes, we compute the gyration tensor and its eigen-values *λ*_1_≤ *λ*_2_ ≤*λ*_3_. To quantify the effective number of spatial directions occupied by a polysome conformation, we introduce the eigenvalue-based shape dimensionality *d*_shape_:

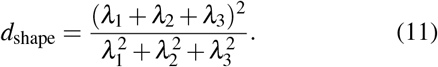

It takes the values 1, 2, and 3 for rod-like, planar, and spherically symmetric conformations, respectively. In three dimensions, this quantity is related to the relative shape anisotropy *κ*^2^ by *d*_shape_ = 3/(1 + 2*κ*^2^)^27^. To further resolve the nature of the shape anisotropy, whether it is prolate or oblate, we also introduce the prolateness Σ^28,29^, defined as

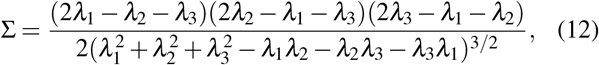

which ranges from − 1 ≤Σ ≤1. It takes the limiting values of 1 for a fully oblate object (such as a flat disk) and −1 for a fully prolate object (such as a rigid rod).

As shown in Fig. 5(left) with *θ* = *π*/2, for linear polysomes, *d*_shape_ and Σ as functions of *N*_space_ behave essentially identically for both ghost and real ribosomes. The shape dimensionality *d*_shape_ decreases from ≈ 1.7 to ≈ 1.4 with decreasing *N*_space_. Simultaneously, the prolateness Σ increases from ≈ 0.75, which is a typical value for an unconstrained flexible chain^18,26^, and approaches 1. This structural reduction toward a quasi-one-dimensional state indicates that the mRNA backbone stretches axially into an elongated, rod-like conformation.

**FIG. 5.**
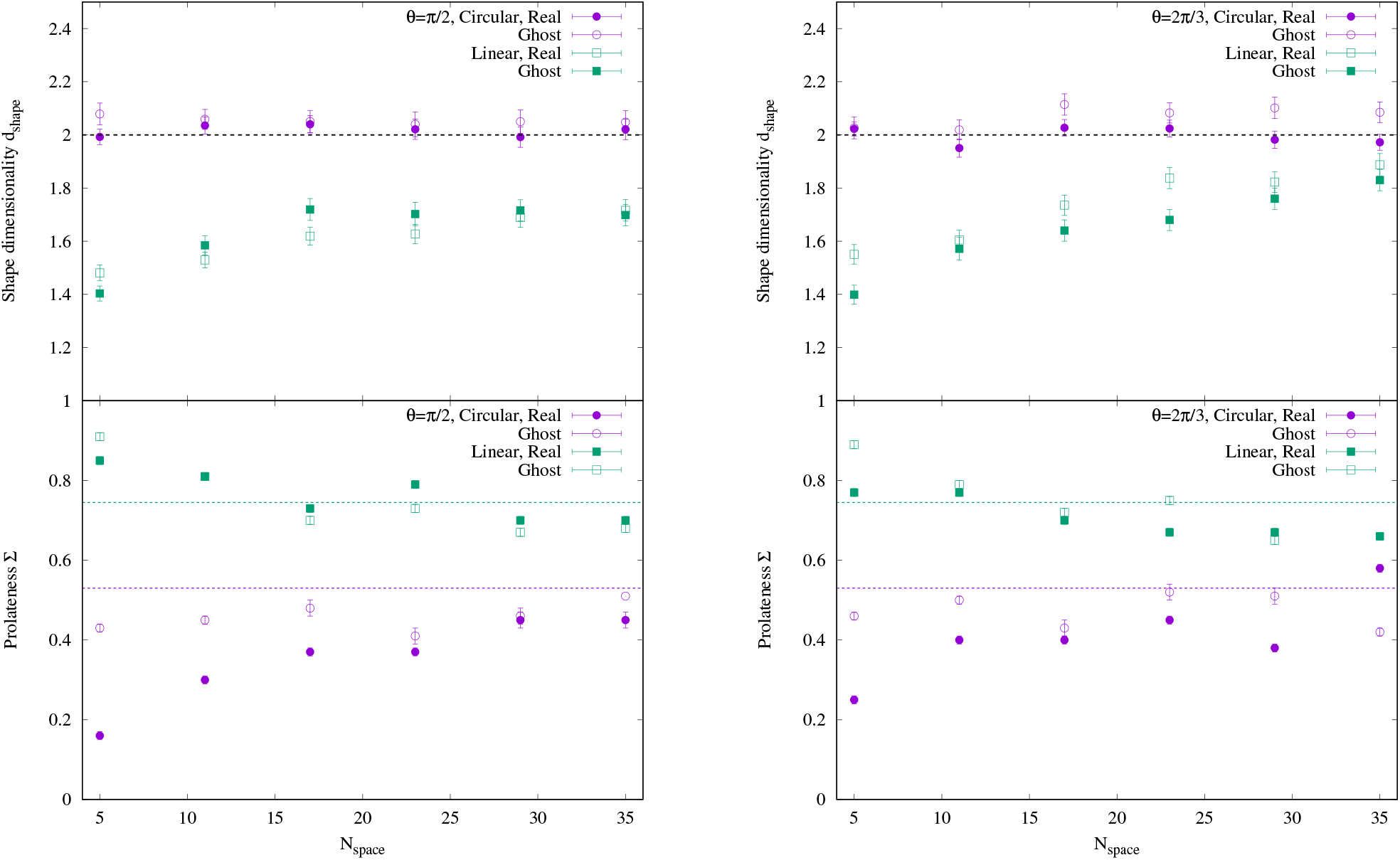
Shape dimensionality *d*_shape_ and prolateness Σ as functions of ribosome spacing *N*_space_ for monomer number *N* = 4969. In the Σ panel, the upper and lower horizontal lines indicate 0.75 and 0.53, which are typical values for flexible linear and circular chains, respectively^18,26^. Symbols represent ghost linear (open squares), real linear (closed squares), ghost circular (open circles), and real circular (closed circles) polysomes. The error bars indicate the standard error across 32 independent replicas. Left: *θ* = *π*/2. Right: *θ* = 2*π*/3.

The comparable macroscopic shape anisotropy between ghost and real ribosomes implies an interplay of physical constraints imposed by the ribosomes. In the ghost model, the increased fraction of rigid ribosome footprints at smaller *N*_space_ naturally stiffens the overall mRNA backbone. Because linear chains have open terminal boundaries, the stiffened chain can freely uncoil and release stress axially. For real ribosomes, the dense steric corridor introduces two additional, competing interactions. On one hand, the steric corridor effect strongly promotes axial swelling. On the other hand, ribosomal excluded-volume interactions geometrically frustrate the chain, preventing the mRNA from adopting a perfectly straight configuration. Consequently, while the whole-chain length increases, local steric bulk also increases the chain thickness (*λ*_1_ and *λ*_2_) compared to the ghost model. To confirm this, the quantitative values at *N*_space_ = 5 are *λ*_1_ = 564.0 ±272.8 and *λ*_2_ = 1909.1 ±851.1 for real ribosomes, compared to *λ*_1_ = 266.3 ±120.2 and *λ*_2_ = 839.7 ±440.0 for ghost ribosomes. These competing local phenomena balance out, preserving a globally elongated, prolate geometry dictated by the open boundaries.

In contrast, for circular polysomes the steric corridor effect cannot escape through terminal boundaries and is uniformly distributed along the mRNA backbone. Crucially, the shape dimensionality of circular polysomes remains constant at *d*_shape_ ≈ 2 across all *N*_space_ for both ghost and real ribosomes. This invariance demonstrates that the closed topology robustly confines the polysome into prolate-like flattened shape, that is a quasi-two-dimensional planar conformation, effectively suppressing out-of-plane thermal fluctuations.

Although the circular polysome is confined to this quasi planar conformation, its in-plane spatial organization undergoes a significant geometric transformation as ribosome density increases. For circular polysomes with real ribosomes, Σ drops from 0.45 to 0.16 with decreasing *N*_space_. In contrast, Σ for ghost ribosomes remains almost constant around 0.53, which is the typical value for a flexible ring polymer (as shown in the bottom panels of Fig. 5).

This reduction in Σ for real ribosomes captures the inplane symmetrization of the polysome ring. In the lowdensity regime (*N*_space_ = 35), the circular backbone forms an anisotropic elliptical shape. However, as *N*_space_ decreases, the steric corridor effect, which has no available escape routes either along the ring contour (due to the loop closure) or in the out-of-plane direction (due to the 2D confinement), generates an outward expansion force along the minor axis of the confined ellipse. As a result, the steric corridor drives the conformation toward a more isotropic, less elongated ring architecture. The invariance of Σ for ghost ribosomes indicates that the effectively longer persistence length never generates a force that expands the minor axis of the confined ellipse, and the planar shape remains unchanged. For *θ* = 2*π*/3, both linear and circular polysomes with ghost and real ribosomes exhibit essentially the same qualitative behavior as for *θ* = *π*/2, as shown in Fig. 5(right), although with some quantitative differences. For linear polysomes, *d*_shape_ is comparably larger and Σ is smaller across the entire *N*_space_ range for both ghost and real ribosomes. For circular polysomes, the qualitative behavior is similar to that for *θ* = *π*/2 across the entire range. Quantitatively, Σ at large *N*_space_ with real ribosomes is ≈ 0.25, which is clearly larger than ≈ 0.16 for *θ* = *π*/2, reflecting a weaker steric corridor effect due to the increased persistence length. This results emphasize that, in all tackled cases, circular polysomes remain robustly confined to a quasi-two-dimensional planar conformation, as evidenced by the invariant *d*_shape_ ≈ 2.

### D. Phenomenological crossover model for the enhancement of scaling exponent *ν*

Here, the terms “three-dimensional” and “quasi-two-dimensional” refer to the operational scaling regimes inferred from *ν*, rather than directly to the finite-size shape dimensionality *d*_shape_. Based on the observed behavior of *ν*, we propose a Phenomenological crossover model describing the crossover between two scaling regimes as a function of the inter-ribosome spacing *N*_space_. In the low-density limit, the global size exponent of the circular polysome approaches the three-dimensional SAW value, *ν*_3D_ ≈ 0.59. As *N*_space_ decreases below a characteristic threshold *N*_*c*_, steric interactions progressively reorganize conformations within the closure-selected quasi-planar ensemble, and the global size exponent approaches a value close to the two-dimensional SAW limit, *ν*_*s*_ ≃ 0.75 (as shown in the bottom panels of Fig. 5). The crossover is therefore a density-dependent change in conformational statistics within an already quasi-planar global geometry, rather than the onset of planarity itself.

Taking into account the smooth crossover broadened by thermal fluctuations, we describe *ν* as a continuous function of *N*_space_:

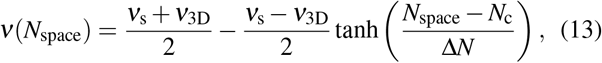

where Δ*N* parameterizes the crossover width of the *ν* enhancement. Because the crossover is governed by the structural fluctuations of the intervening flexible mRNA subchain (excluding the bound ribosome footprint of length 30 monomers), the subchain size scales as 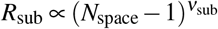. Consequently, Δ*N* is derived from the standard deviation of *R*_sub_ around *N*_c_ as:

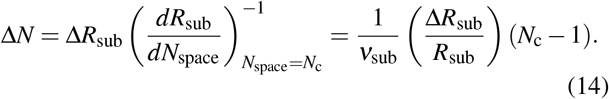

While the relative fluctuation of a polymer subchain Δ*R*_sub_/*R*_sub_ is known to be around 0.40 in standard polymer statistics, i.e. 0.42 for ideal chain and 0.36 for the self avoiding chain^30^, we consider that Δ*R*_sub_/*R*_sub_ in our system takes a smaller value, reflecting the highly suppressed fluctuation and stabilized architecture of the circular polysome. By directly calculating *ν*_sub_ and Δ*R*_sub_/*R*_sub_ from our simulations, Eq. (14) establishes that the transition width Δ*N* is naturally proportional to *N*_c_. In this formulation, *ν*_sub_ and Δ*R*_sub_/*R*_sub_ are independently determined from the subchain structure factor and the end-to-end distance fluctuation, respectively, and are used as inputs to Eq. (14) to predict the crossover width Δ*N*, rather than treating it as a free fitting parameter. Consequently, the nonlinear fit to Eq. (13) involves only two adjustable parameters: the saturated exponent *ν*_s_ and the threshold spacing *N*_c_.

To determine the subchain scaling exponent *ν*_sub_ required for our model and to validate the global scaling behavior in reciprocal space, we calculate the structure factor *S*(*q*) for the circular polysomes:

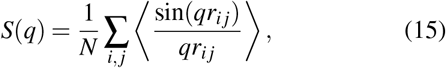

where *q* is the wave vector and *r*_*i j*_ is the distance between the *i*-th and *j*-th monomers. Note that only the mRNA backbone monomers are considered in this calculation.

According to standard polymer scaling theory^31,32^, the structure factor follows a power-law decay *S*(*q*) ∝ *q*^−1/*ν*^ within the fractal internal scaling regime, strictly defined by *qR*_g_ ≫ 1. Practically, a previous study analyzing semi-flexible cyclic macromolecules^33^ demonstrated via generalized Kratky plots that the internal scaling regime (exhibiting either a flexible plateau or semi-flexible oscillatory behavior) robustly emerges at *qR*_g_ ≳ 4. Hence, for our circular polysome systems, we define the window *q* > 4.3/*R*_g_ as the accessible internal scaling regime.

We plot *S*(*q*) for *N* = 4969 with *θ* = *π*/2 and 2*π*/3 across *N*_space_ = 5, 11, and 17 in Fig. 6. Although the curves in Fig. 6 (left) are vertically shifted for visual clarity, the unscaled *S*(*q*) curves nearly overlap in the high-*q* region (*q* > 0.2*u*^−1^) while exhibiting a small plateau in the low-*q* region (*q* < 0.1*u*^−1^). The position of this plateau shifts depending on *N*_space_. Specifically, for *N*_space_ = 5 (*R*_g_ = 88.1*u*, calculated considering mRNA backbone), the plateau appears around *q* ≈ 0.04*u*^−1^, which closely matches *π*/*R*_g_. This indicates that this feature corresponds to the global diameter of the circular polysome.

**FIG. 6.**
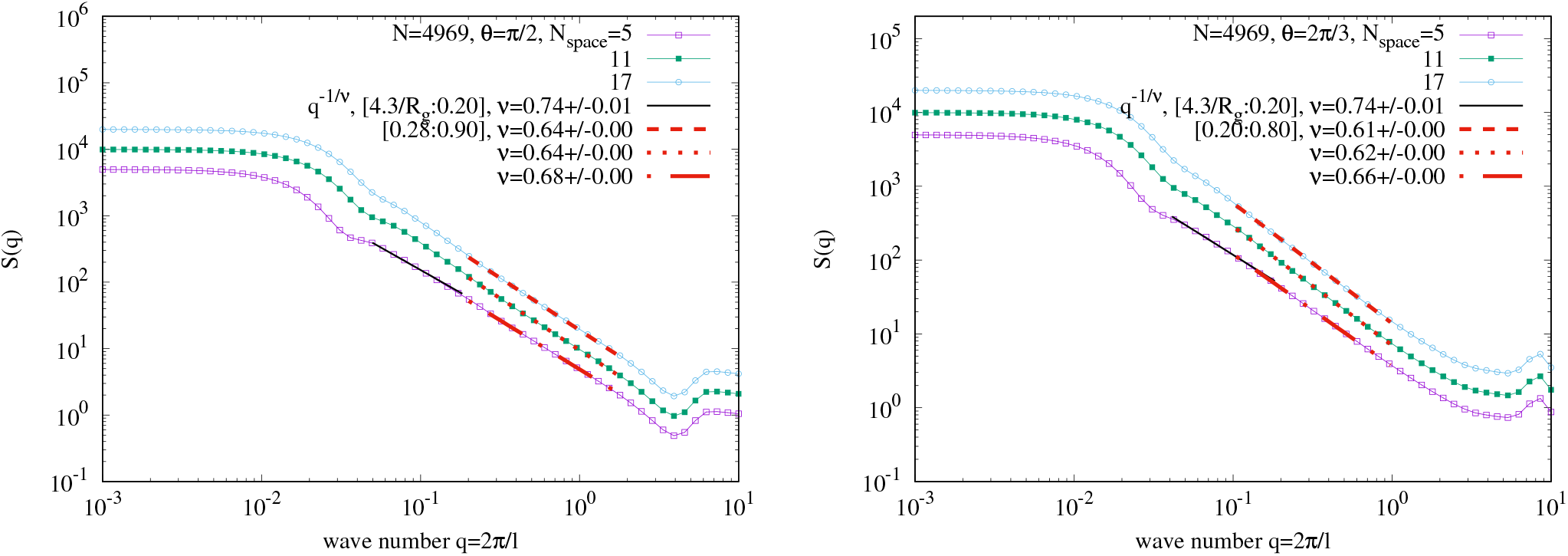
Structure factor *S*(*q*) as a function of wavenumber *q* for monomer number *N* = 4969. The curves are vertically shifted for clarity and represent *N*_space_ = 5, 11, and 17, from bottom to top. The lines indicate least-squares fits to *S*(*q*) ∝ *q*^−1/*ν*^, where dashed, dotted, and dot-dashed lines correspond to *N*_space_ = 17, 11, and 5, respectively. Left: *θ* = *π*/2. Right: *θ* = 2*π*/3.

To avoid interference from this global shape factor, we fit the structure factor within the window 4.3/*R*_g_ < *q* < 0.2*u*^−1^. For *N*_space_ = 5, the fit of *q*^−1/*ν*^ yields *ν* = 0.74 ± 0.01. Crucially, this value is highly consistent with the scaling exponent obtained from 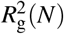 in real space within twice the standard error. For *N*_space_ = 11 and 17, a stable power-law fit in this global regime cannot be isolated due to the narrowing of the wave-number window between 4.3/*R*_g_ and the subchain threshold *q*_sub_.

Beyond *q* ≈ 0.2*u*^−1^, the scaling behavior deviates from this global exponent. Fitting the intermediate *q*-range of 0.28*u*^−1^ < *q* < 0.9*u*^−1^ yields a different set of exponents: *ν* = 0.68, 0.64, and 0.64 for *N*_space_ = 5, 11, and 17, respectively. In real space, this wave number range corresponds directly to the length scale of the subchains between adjacent ribosomes, where the subchain *R*_g_ is 11.0*u* for *N*_space_ = 5, 17.9*u* for *N*_space_ = 11, and 22.7*u* for *N*_space_ = 17. We confirmed that varying the fitting window by ± 10% alters *ν*_sub_ by less than 2%, which introduces negligible error into the determination of *N*_c_. Hence, we estimate the subchain scaling exponent *ν*_sub_ for any given density via linear interpolation of these values for *θ* = *π*/2.

The structure factor for *θ* = 2*π*/3 (Fig. 6, right) exhibits qualitatively identical behavior. For the highest density case (*N*_space_ = 5), the fit in the large-scale window (4.3/*R*_g_ < *q* < 0.2*u*^−1^) yields *ν* = 0.74 ± 0.01, again in excellent agreement with the real-space 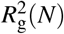 analysis. The fit in the subchain window (0.2*u*^−1^ < *q* < 0.8*u*^−1^) for *N*_space_ = 5, 11, and 17 yields *ν* = 0.66, 0.62, and 0.61, respectively. Thus, we similarly estimate *ν*_sub_ by linear interpolation of these values for *θ* = 2*π*/3.

To evaluate the relative fluctuation of the mRNA subchains, Δ*R*_sub_/*R*_sub_, we directly calculate the standard deviation and mean of the end-to-end distance for subchains between adjacent ribosomes from our simulations across *N*_space_ = 5 to 35 for both *θ* = *π*/2 and 2*π*/3, and plot them in Fig. 7.

**FIG. 7.**
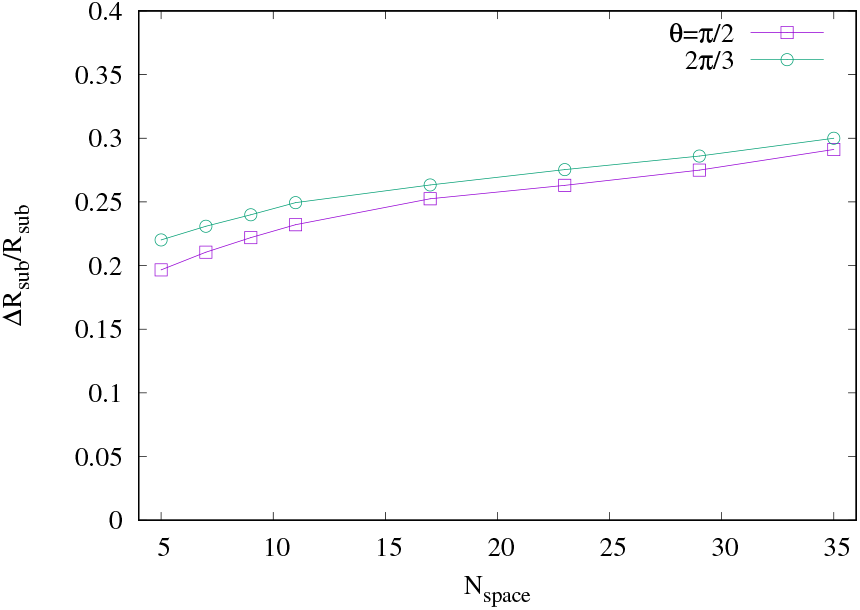
Relative fluctuation of the subchain end-to-end distance, Δ*R*_sub_/*R*_sub_, as a function of ribosome spacing *N*_space_ for monomer number *N* = 4969, where *R*_sub_ is the end-to-end distance of the subchain between adjacent ribosomes and Δ*R*_sub_ is the standard deviation of it. Symbols represent local bending angles *θ* = *π*/2 (open squares) and *θ* = 2*π*/3 (open circles).

As *N*_space_ decreases, the relative fluctuation decreases from around 0.30 to substantially lower values. Furthermore, the fluctuation for *θ* = *π*/2 is systematically smaller than that for *θ* = 2*π*/3. This indicates that a shorter subchain distance, resulting from a shorter persistence length, imposes stronger steric constraints and further suppresses conformational freedom. All observed values remain distinctly below the benchmark ratio of ≈ 0.40 typical for unconstrained flexible chains^30^. This reduction in fluctuation confirms that circular polysomes form a structurally stabilized architecture whose rigidity increases with ribosome density, fully consistent with the accelerated relaxation dynamics observed in the auto-correlation analysis of 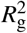. In our analysis, these numerical data are continuously interpolated to dynamically evaluate Δ*R*_sub_/*R*_sub_ at *N*_space_ = *N*_c_ during optimization. Incorporating this into Eq. (14) uniquely computes the transition width Δ*N* at each iteration, leaving only two free fitting parameters (*ν*_s_ and *N*_c_) for the crossover model.

Based on the parameters determined above, we fit the crossover model described by Eq. 13 using *ν*_s_ and *N*_c_ as free parameters via the non-linear least-squares method. As shown in Fig. 8, the fitting results show excellent agreement with the values of *ν* obtained from our simulations within error bars. This quantitative agreement demonstrates that our two-parameter Phenomenological crossover model captures the essential mechanism underlying the density-dependent enhancement of *ν*.

**FIG. 8.**
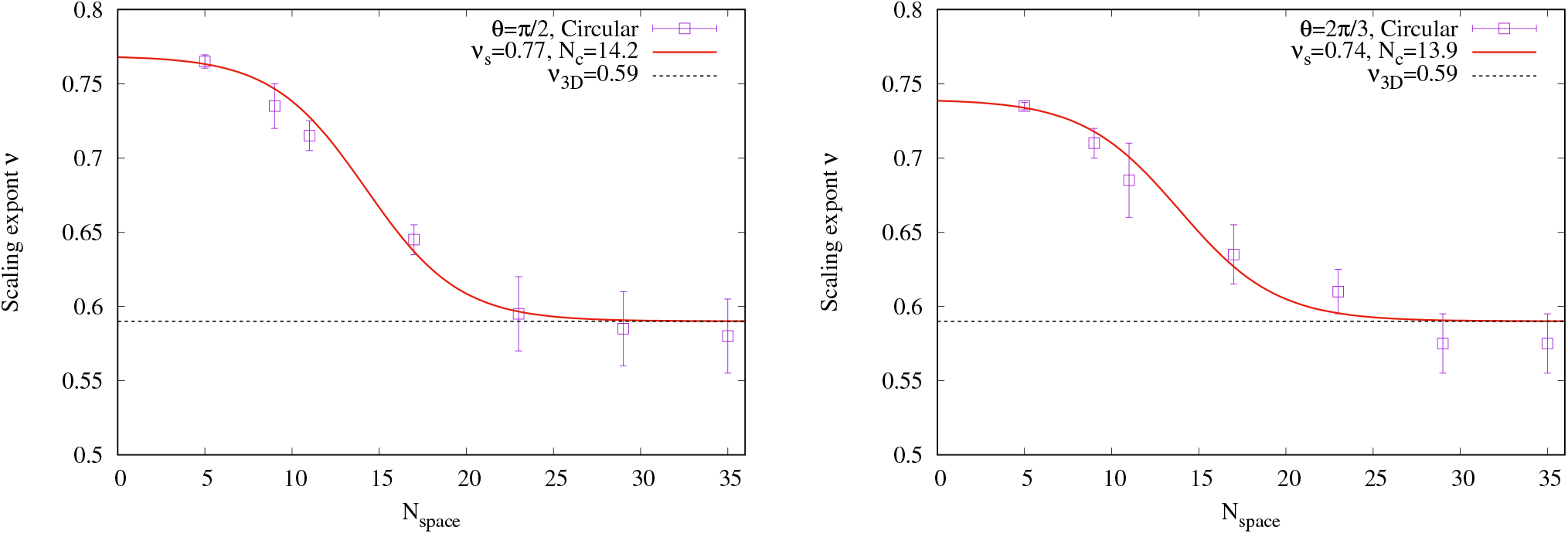
Scaling exponent *ν* obtained from 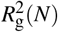 as a function of ribosome spacing *N*_space_. The error bars indicate fitting uncertainties for *ν*. The solid lines indicate least-squares fits to Eq. (13). Left: *θ* = *π*/2. Right: *θ* = 2*π*/3.

The saturated scaling exponent yields *ν*_s_ = 0.77 ± 0.01 for *θ* = *π*/2, which is slightly larger than *ν*_s_ = 0.74 ± 0.01 for *θ* = 2*π*/3. This difference suggests that a shorter persistence length (and thus a tighter local bending geometry) imposes stronger steric constraints along the mRNA backbone. Applying the Flory-type relation, the corresponding effective spatial dimensions are evaluated as *d*_eff_ = 1.90 for *θ* = *π*/2 and *d*_eff_ = 2.05 for *θ* = 2*π*/3. Both values correspond, within the Flory-type operational interpretation, to an effective dimension close to two. They indicate that the dense-limit scaling behavior is consistent with two-dimensional-SAW-like statistics, regardless of the bending angle considered here.

Furthermore, the characteristic threshold spacing is estimated as *N*_c_ = 14.2 ±0.4 for *θ* = *π*/2 and *N*_c_ = 13.9 ±1.1 for *θ* = 2*π*/3. These values show no significant difference within fitting uncertainties. This angular invariance of *N*_c_ can be understood by comparing the subchain length with its conformational fluctuations. At the subchain level, the mean end-to-end distance *R*_sub_ for *θ* = 2*π*/3 is only ≈ 10% longer than that for *θ* = *π*/2. This difference is substantially smaller than the intrinsic thermal fluctuations of the subchains (Δ*R*_sub_/*R*_sub_ ≳ 20%). Because thermal fluctuations broaden the spatial distributions and make the spatial extent of the subchains physically comparable across bending angles, the onset threshold for the steric corridor remains essentially constant around *N*_c_ ≈ 14. This invariance indicate that the onset of the steric corridor is a universal geometrical threshold governed by ribosome spacing rather than local chain stiffness.

### E. Relative fluctuation

Finally, to evaluate how structural stability depends on ribosome density (*N*_space_), as well as the interplay between chain topology and steric interaction, we calculate the relative fluctuation of the radius of gyration, Δ*R*_g_/*R*_g_, where Δ*R*_g_ represents the standard deviation of *R*_g_. We compare both linear and circular polysomes using two ribosome models: “ghost” ribosomes (without steric excluded volume) and “real” ribosomes [Fig. 9].

**FIG. 9.**
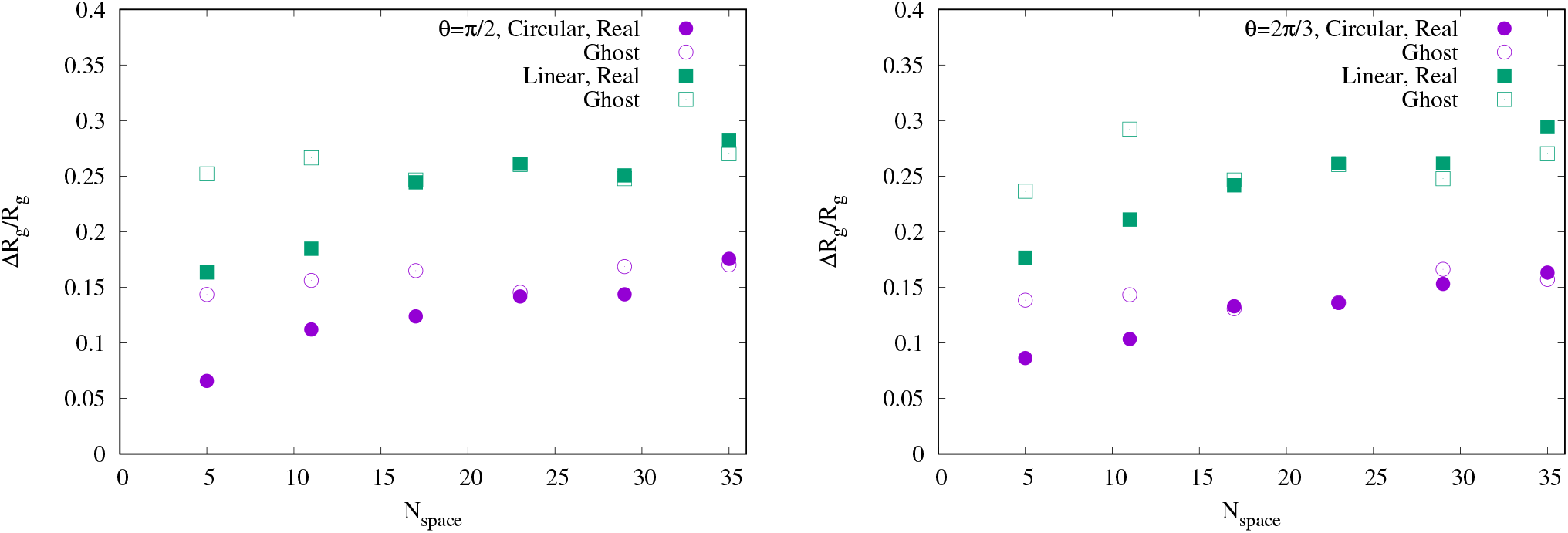
Relative fluctuation of the radius of gyration, Δ*R*_g_/*R*_g_, as a function of ribosome spacing *N*_space_ for monomer number *N* = 4969, where Δ*R*_g_ is the standard deviation of *R*_g_. Symbols represent ghost linear (open squares), real linear (closed squares), ghost circular (open circles), and real circular (closed circles) polysomes. Left: *θ* = *π*/2. Right: *θ* = 2*π*/3.

For *θ* = *π*/2, the ghost ribosome systems shows relative fluctuations that are essentially independent of *N*_space_ [Fig. 9(left)]. Specifically, the baseline fluctuation for linear polysomes with ghost ribosomes (≈ 0.26) is distinctly larger than that for circular polysomes (≈ 0.16). This demonstrates that the circular topology stabilizes the global architecture even in the absence of steric corridor constraints. In contrast, for real ribosomes, the relative fluctuation decreases systematically with decreasing *N*_space_ once the density exceeds the characteristic threshold (*N*_space_ < *N*_c_). Across the ribosome density range up to *N*_space_ = 5, the steric crowding yields a comparable absolute reduction in the relative fluctuation amplitude for both topologies, Δ(Δ*R*_g_/ ⟨ *R*_g_ ⟩) ≈ 0.09–0.10. Specifically, the relative fluctuation drops from 0.26 to 0.16 (38% reduction) for linear polysomes, and from 0.16 to 0.07 (56% reduction) for circular polysomes. Interestingly, the fluctuation level reached by the most densely loaded linear polysome (≈ 0.16) merely matches that of a passive circular ring with ghost ribosomes (≈ 0.16). Because topological closure operates from this already reduced baseline, the comparable steric-driven decrease pushes the circular system into an exceptionally stable state of 0.07, which is less than half that of the dense linear counterpart. This comparison confirms that the enhancement of conformational stability in circular polysomes arises from a non-additive, synergistic coupling between topological closure and steric constraints, driving the system into a high-stability regime unattainable by open linear architectures.

Systems with *θ* = 2*π*/3 display qualitatively identical behavior [Fig. 9(right)]. For ghost ribosomes, the relative fluctuations remain *N*_space_-independent at 0.26 for linear polysomes and 0.15 for circular polysomes. For real ribosomes, the relative fluctuations once again decrease continuously below *N*_c_ as *N*_space_ decreases. These findings show that closure and steric interactions act synergistically to suppress global fluctuations, while playing distinct roles in establishing the global geometry and its density-dependent in-plane reorganization.

## IV. DISCUSSION

The central finding of this work is that mRNA circularization fundamentally alters polysomes’ conformational response to ribosomal crowding. Circular closure and ribosome-mediated steric interactions do not act as two independent, additive constraints on the mRNA backbone. Instead, their interplay establishes a distinct conformational state characterized by strongly suppressed global-size fluctuations. This closure-crowding interplay is most directly demonstrated by the four-system comparison in Fig. 9. In the absence of inter-ribosomal excluded volume, closure alone reduces the relative gyration-radius fluctuation from Δ*R*_g_/*R*_g_ ≃ 0.26 for linear polysomes to ≃ 0.16 for circular polysomes. Steric interactions similarly reduce the fluctuation of linear polysomes to ≈ 0.16 at maximum ribosome density. However, the combination of circular closure and steric crowding cooperatively suppresses the fluctuation down to Δ*R*_g_/*R*_g_ ≃ 0.07. Thus, the densely ribosomes loaded circular polysome accesses a conformationally stabilized state unattainable by closure alone or by steric crowding in an open architecture.

The physical essence of this state lies in the topology dependence of the crowding response, rather than a simple additive decomposition of fluctuations. In fractional terms, increasing the ribosome density reduces Δ*R*_g_/*R*_g_ by approximately 38% in the linear system, whereas the corresponding reduction in the circular system reaches 56%. This difference is crucial because ring closure decreases the initial fluctuation level. A larger percentage reduction in an already constrained state indicates that closure and crowding act together to suppress fluctuations. The ghost-ribosome controls are essential to this conclusion, as increasing the footprint-fraction without inter-ribosomal excluded volume fails to reproduce the density-dependent stabilization observed for circular polysomes with real ribosomes. In other words, the observed stabilization cannot be attributed to the increased local stiffness imparted by ribosome-bound mRNA footprints. The structural observables reveal how this stabilized state is organized. Circular closure selects a global conformational ensemble with an approximately density-independent shape dimensionality of *d*_shape_ ≈ 2 for *N* = 4969. This quasi-planar geometry separates the topological consequence of closure from the density-dependent steric response. Within this closure-selected ensemble, real ribosomes drive pronounced in-plane reorganization: as ribosome density increases, prolateness decreases as the circular polysome transitions from an elliptical to a more isotropic, circularized conformation. Because ghost-ribosome systems lack this systematic shape evolution, this structural response is not a passive consequence of closing a semiflexible chain, but requires cooperative steric interactions among the ribosome-decorated segments across the closed backbone.

The topology-dependent response is also obvious in the global size scaling. At low ribosome density (*N*_space_ ≥ 23), the scaling exponent *ν* of circular polysomes converges to the standard three-dimensional self-avoiding walk (3D-SAW) value (*ν* ≈ 0.588). At high loading density (*N*_space_ = 5), *ν* increases toward 0.74–0.77 depending on the local bending angle *θ* . We interpret this enhancement as a density-induced crossover toward a swollen, quasi-two-dimensional scaling regime within the closure-selected ensemble. This crossover must be distinguished from a second onset of planarity, as *d*_shape_ ≈ 2 remains constant across all densities. Instead, steric crowding reorganizes the conformational statistics within the quasi-planar architecture by suppressing the fluctuations that permit a low-density ring to remain elliptical.

The scale separation between local subchains and global topology explains why the onset threshold *N*_c_ ≈ 14 is robust and virtually angle-insensitive, whereas the saturated exponent *ν*_s_ exhibits angle dependence. At the individual subchain scale, the modest ≈ 10% difference in average length induced by local bending angles (*θ*) is masked by strong thermal fluctuations (Δ*R*_sub_/*R*_sub_ ≳ 20%). At the global chain scale, however, topological closure prevents steric stress from escaping through open ends. Under high-density crowding, extending a subchain forces adjacent ribosomes into closer spatial proximity. For *θ* = *π*/2, this steric restriction is inherently more severe. As a result, local packing constraints propagate and accumulate cooperatively across the entire closed loop, yielding a higher saturated exponent (*ν*_s_ ≈ 0.77 vs 0.74). The fluctuation-informed crossover model captures this scale-dependent coupling by directly linking the crossover width to the measured size fluctuations of inter-ribosomal subchains.

The intermediate-*q* exponent of circular polysomes decouples from the global scaling behavior rather than remaining scale-invariant. In the crowded regime for *θ* = *π*/2, the global exponent is enhanced toward *ν* ≃ 0.76, whereas the subchain exponent remains lower (*ν*_sub_ ≃ 0.68 for *N*_space_ = 5), closer to the 3D-SAW value. Once ribosome density decreases such that the spacing exceeds the crossover threshold (*N*_space_ = 17 > *N*_*c*_), the global exponent decreases to the subchain value, and this separation vanishes within fitting uncertainties. In contrast, linear polysomes show an essentially scale-invariant exponent across the accessible *q*-range, consistent with a self-similar swelling of the steric corridor along the open backbone^19^. The exponent separation in the circular case is thus consistent with a crowded, cooperatively swollen regime that promotes a 2D-SAW-like global expansion. Because topological closure is a global constraint, its direct effect on subchains between adjacent ribosomes is naturally suppressed. This agrees with our two-scale crossover model, where subchain statistics enter only into Δ*N* without altering *ν*_*s*_. We note that, for linear polysomes, the *S*(*q*) fit was restricted to 0.5 *u*^−1^ < *q* < 1.0 *u*^−1^ due to low-*q* noise at lower *q* in the present replica statistics. An open question for circular polysomes is how global topological closure drives this scale-dependent exponent deviation. Future studies will focus on its microscopic origin by incorporating experimental data from both cryo-ET and smFISH measurements, which provide also ribosome orientations and effective gyration radius, respectively.

Closure also alters the internal relaxation dynamics of the crowded chain. In linear polysomes, increasing ribosome density broadens the relaxation spectrum of *R*_g_ fluctuations and amplifies their variability, without systematically accelerating the persistently slow decorrelation time. In circular polysomes, a systematic opposing trend emerges. The stretching exponent increases toward unity, and the decorrelation time progressively decreases with crowding. At the highest ribosome density, global size decorrelation is approximately 45 times faster in the circular system than in its linear counterpart. Although our minimal polymer model omits hydrodynamic interactions and active kinetics, this stark contrast isolates a purely topological mechanism, as static stabilization and rapid conformational relaxation are fully compatible between both linear and circular CG polysomes. While the dense circular polysome possesses a severely restricted fluctuation amplitude, its residual collective fluctuations relax rapidly with a narrow distribution of relaxation times, protecting the closed architecture from steric-induced kinetic arrest.

Comparing circular with linear polysomes represents the fundamental role of boundary conditions in overcrowded biopolymers. In open architectures, steric constraints along the backbone can be accommodated through conformational rearrangements and stress release at the chain terminals. Circular closure eliminates this boundary-mediated relaxation pathway, forcing steric interactions to distribute cooperatively across the entire contour. The resulting state is neither a simple extrapolation of the linear steric corridor nor a passive property of an isolated polymer ring, but a topology-dependent, collective reorganization of a crowded macro-molecular assembly.

While our study focuses on equilibrium conformational statistics and internal relaxation dynamics rather than direct translational kinetics, these findings imply that circularizing mRNA fundamentally reshapes the mechanical and steric environment experienced by translating ribosomes. The simultaneous suppression of global size fluctuations and preservation of rapid internal decorrelation may optimize polysome spatial organization in the crowded cytoplasm.

There are several biological features limiting the quantitative comparison between experiments and our multi-polysome model that will be explicitly tackled in our future research, namely: hydrodynamic interactions^34,35^, ribosome-ribosome coordination^36–38^ and active translocation^39–41^. Among other higher resolution features e.g. codon dependent stepping^42–44^ or effects from secondary structures formation^45–48^, that will be tackled by enhancing the multiscale character of our model to specific bases in a top-down modeling strategy.

## V. CONCLUSION

In this study, we distinguished the geometric consequence of circular closure from the density-dependent steric response of ribosome-decorated mRNA. Circular architecture selects a quasi-planar global conformational ensemble (*d*_shape_ ≈ 2), which remains virtually invariant across all examined ribosome loading densities. Ribosome crowding does not induce this planar character; rather, inter-ribosomal steric interactions drive a cooperative structural reorganization within this closure-selected ensemble. With increasing ribosome density, circular polysomes transition from an elliptical to a more planar isotropic conformation, their global size scaling exponent increases toward *ν*_s_ ≈ 0.74–0.77, and their relative size fluctuations drop to Δ*R*_g_/*R*_g_ ≈ 0.07. The fluctuation-informed crossover model quantitatively connects this global scaling to subchain-scale statistics, explaining how local thermal fluctuations preserve a robust, angle-insensitive onset threshold (*N*_c_ ≈ 14), whereas global topological constraints amplify cumulative steric stress across the closed backbone.

The same closure-crowding interplay fundamentally transforms global-size relaxation dynamics. Whereas dense linear polysomes exhibit slow, heterogeneous *R*_g_ relaxation with a broadened spectrum, dense circular polysomes display an accelerated decorrelation time approximately 48 times faster than matched linear systems, accompanied by a markedly narrowed relaxation spectrum. Thus, circularization converts the crowding response into a structurally stabilized state without dynamic arrest. From a biophysical perspective, these find-Topological Closure Drives Structural Stabilization and Fast Cooperative Dynamics in Crowded Circular Polysomes ings show that mRNA circularization alters the mechanical and conformational environment experienced by translating ribosomes far beyond merely bringing the 5’ and 3’ ends into spatial proximity. Determining whether this structurally stabilized yet fast-relaxing conformational state modulates ribosome recycling, traffic congestion, or overall translation efficiency is an important direction for future multi-polysome models that explicitly account for active forces and hydrodynamic interactions.

## ACKNOWLEDGMENTS

We thank Brian Zid for illuminating discussions on the experimental measurements of mRNA during translation, Anton Petrov for sharing important literature of polysomes in different cellular contents and Mahesh Yadav for discussions on graphical polysome representations. H.K. thanks Gemini 3.7 for its assistance with grammatical correction and prose flow, both of which were subsequently reviewed and approved by the authors. H.V.G. acknowledges financial support from the Ramón y Cajal grant No. RYC2022-038082-I and Spanish Ministry of Science and Innovation, through project PID2023-150536NA-I00, and the “Severo Ochoa” Grant No. CEX2023-001263-S for Centers of Excellence; and CSIC’s grant MMT24-ICMAB-01 for the nanoML4Med project. H.V.G. acknowledges also Red Española de Supercomputación (RES) for the computing time and technical support at the Marenostrum 5 FI-2026-1-0062. HK acknowledge funding from the Deutsche Forschungsgemeinschaft(DFG, German Research Foundation) under grant No 528726435. The authors dedicate this work to the memory of Prof. Rudi Podgornik, and H.V.G. is grateful for years of fruitful collaborations and scientific discussions, in particular on the topic of developing methods for multiscale simulations of RNA, exploiting polymer concepts in biological systems.

